# An automated approach for systematic detection of Key Biodiversity Areas

**DOI:** 10.1101/2023.06.30.547173

**Authors:** Dario Nania, Gentile Francesco Ficetola, Mattia Falaschi, Michela Pacifici, Maria Lumbierres, Carlo Rondinini

## Abstract

The new Key Biodiversity Areas (KBA) standard is an important method for identifying regions of the planet hosting unique biodiversity. KBAs are identified through the implementation of threshold-based criteria that can be applied to any target species and region. Efficient methods to rapidly assess the existence of potential KBAs in different areas of the planet are still missing, although they are needed to accelerate the KBA identification process for large numbers of species globally. We developed a methodology to scan geographical regions and detect potential KBAs under multiple criteria. We tested the methodology on 59 species of reptiles and amphibians in Italy through the application of selected KBA criteria. Potential KBAs were identified for multiple species under most criteria, covering 1.4% to 12% of the study area, depending on analytical settings. Unit size used to identify KBAs played an important role in shaping the distribution of potential KBAs, also affecting the degree of overlap between areas triggered by different criteria. New potential KBAs identified in this study are only partially nested within current KBAs in Italy (previously identified for birds) and within the national protected areas.

## Introduction

Global change is profoundly affecting Earth’s biodiversity (IPBES, 2019). As species decline and their extinction risk continues to grow globally (Hoffmann et al., 2010), there is an urgent need to identify sites that are important for biodiversity persistence. Currently, there are several methods to identify important sites for biodiversity, some of them being target based and others threshold based (Smith et al., 2018). The most recent Key Biodiversity Areas (KBA) standard was developed to identify sites that can contribute to the global persistence of biodiversity (IUCN, 2016).

The KBA approach has been included by the Convention on Biological Diversity as a valuable resource to inform protected areas expansion for the achievement of the Aichi Biodiversity Targets (CBD, 2022). The Important Bird Areas (IBA), which are now part of the KBA network, have been used by the European Union to design protected areas under its first legislation on the environment, the Birds Directive (Directive 2009/147/EC on the conservation of wild birds). KBAs have been included by the United Nations in the Sustainable Development Goals Report (UN DESA, 2021) to assess the efficiency of protected areas in covering sites that can ensure the persistence of biodiversity. The KBA approach is therefore already being used extensively in conservation policy.

Globally, 79% of currently recognized KBAs have been identified based on the presence of bird species (keybiodiversityareas.org, April 11th 2022), revealing an important gap of knowledge on the distribution of KBAs for non-avian taxa. In order to reduce the taxonomic bias of KBAs, it would be particularly useful to rely on methodologies that can rapidly assess KBAs within large areas for high numbers of species. Potential KBAs have been identified for different taxa, including vertebrates, invertebrates, and plant species (Ambal et al., 2012; Yahi et al., 2012; Plumtree et al., 2019) using different approaches depending on the geographic scale of the analysis, as well as on the target taxa and data availability. However, such methods do not allow simultaneous KBA assessments for large numbers of species, resulting in a significantly slow process of mapping KBAs worldwide.

We developed a methodology that scans a geographic region to detect potential KBAs for target species based on their habitat distribution. We applied our approach to Area of Habitat (AOH) maps. The AOH maps provide information on the spatial distribution of available habitat for the species within its distribution range and altitude limits (Rondinini et al., 2011; Lumbierres et al., 2022). AOH represents a useful tool for large-scale assessments of the geographic distribution of species’ populations for conservation purposes (Brooks et al., 2019), and is considered a valid proxy of population size by the Global Standard for the Identification of Key Biodiversity Areas (IUCN, 2016). The methodology can be used to efficiently apply KBA criteria which rely on population size and distribution of the examined taxa.

We tested the method on Italian reptiles and amphibians, for which KBAs have not been identified so far. Italy is an important center of diversity and endemism of European herpetofauna (Sindaco et al., 2006; Cox & Temple 2009; Sillero et al, 2014). Seven amphibians and two reptiles occurring in Italy are listed as threatened by the Red List of the International Union for Conservation of Nature (IUCN, 2021). The geographical distribution as well as habitat availability for many reptiles and amphibian species in Italy is well known (IUCN, 2020; Nania et al., 2022). We assessed the effects of KBA criteria, grid size, and species attributes (such as distribution extent, degree of endemism and IUCN Red List status) on the total area of potential KBAs identified. We then compared the distribution of new potential KBAs with the current KBA and protected area networks of Italy.

## Methods

### Species distribution maps

We estimated the AOH of 59 species of amphibians and reptiles within their geographic range in Italy. For 55 species, we used AOH maps from Nania et al. (2022), These maps were built using the ranges downloaded from the IUCN Red List of Threatened Species (IUCN 2021). For four species endemic to Italy we built new AOH maps based on ranges that were obtained from specific Italian datasets because of their higher accuracy. In particular:

- For three species endemic to Italy, the range was obtained from the Atlas of Amphibians and Reptiles of Italy, curated by the Societas Herpetologica Italica (SHI). The maps are available upon request from SHI;
- For one species microendemic to Italy (*Podarcis raffonei*), we used the range from Ficetola et al. (2021). Since the range is very small (5000 m^2^), we did not calculate an AOH and considered the whole range suitable.

These four AOH maps were developed following the procedure shown in Nania et al. (2022). Species were mapped to their habitat using Copernicus Global Land Service Land Cover (CGLS-LC100) 2019 classes as a habitat surrogate. We compiled species-habitat associations and altitude limits based on monographs on the biology of target species (Sindaco et al., 2006; Lanza et al., 2007; Corti et al., 2011). We then reclassified a base map combining land cover and altitude information to produce area of habitat maps for the target species. A complete description of how the additional AOH maps were produced is included in Appendix S1.

### Systematic identification of KBAs

To systematically identify candidate KBAs under five selected criteria (A1, B1, B2, B3, E), we produced four grids of squared cells of different size (10×10, 20×20, 30×30 and 40×40 km) in a Lambert-Azimuthal equal area projection (ETRS89, EPSG:3035) in GRASS GIS version 7.8.5 (GRASS 2020). The grids were built on the country’s land surface, covering the Italian peninsula and its islands. Cell size was selected based on the current mean and median area of reptiles and amphibians KBAs worldwide, according to the Key Biodiversity Areas Secretariat database (https://www.keybiodiversityareas.org/). Currently, the global mean size of KBAs triggered by reptile or amphibian species is 1400 km^2^ (SD = 16910), while the median is 240 km^2^. Thus, by selecting 40×40 km as the largest cell size, we ensured that it could potentially include a KBA with a maximum size of 1600 km^2^, which is slightly above the current average KBA size for reptiles and amphibians worldwide. The smallest cell size can detect KBAs with a maximum size of 100 km^2^. The four grid resolutions allowed us to test for sensitivity of the KBA criteria to the cell size, and therefore to the dimension of the unit used to detect KBAs. To avoid missing potential KBAs due to the fixed position of the grid on the country’s surface, for each resolution we replicated the grid in three additional positions. Specifically, the grid was moved to half the size of the grid cell along two cardinal directions (North and East) and one ordinal direction (North-east), as illustrated in Figure 1. Each of the 16 grids (four resolutions by four positions) was used as a basis for assessing the existence of potential KBAs following the procedure described below. The GRASS and R code used for the analysis is available in Appendix S2.

**Figure 1:**
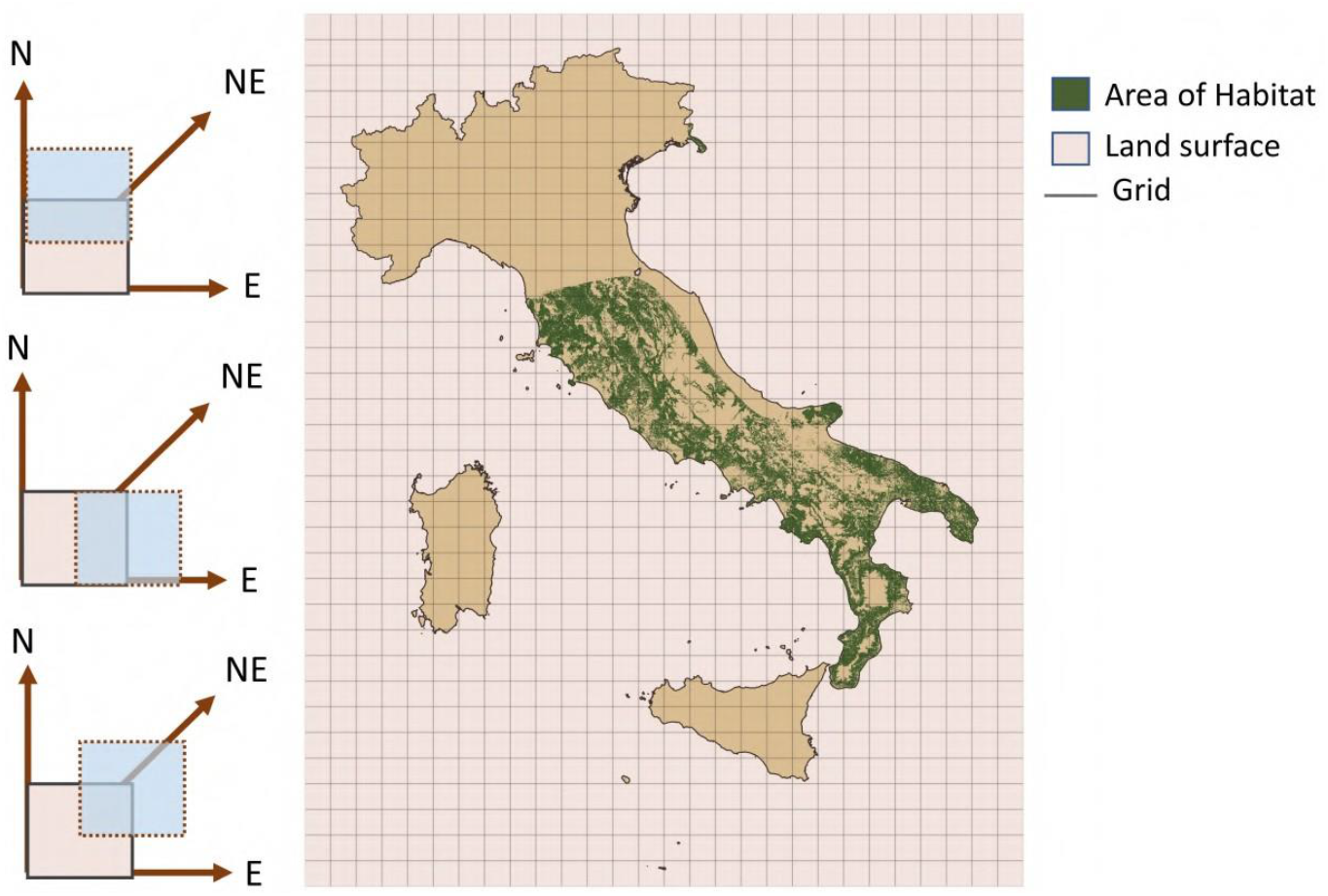
An explanatory scheme showing how the moving window approach works. Here we show the 40×40 km grid positioned on top of the AOH map. During the surface scanning, the grid is moved to three additional positions (left).

Potential KBAs for target species are detected through the application of several threshold-based criteria, which rely on the distribution of important populations of one or more species. The main five KBA criteria refer to “Threatened biodiversity” (criterion A), “Biogeographically restricted biodiversity” (criterion B), “Ecological integrity” (criterion C), “Biological processes” (criterion D) and “Irreplaceability through quantitative analysis” (criterion E) (IUCN 2016). We tested selected criteria/subcriteria that can be informed by the AOH as detailed below.

Criterion A1 refers to globally threatened species. According to the guidelines for KBA identification (IUCN, 2020), a site qualifies as a KBA under criterion A1 if it hosts a minimum threshold percentage of the global population of a species. The percentage threshold depends on the threatened status of the species. For Critically endangered (CR) and endangered (EN) species, the threshold is ≥0.5% of the global population. For a species assessed as vulnerable (VU), the threshold is ≥1%. Species for which criterion A1 was applied are indicated in Table 1. For each species, we calculated the proportion of AOH inside each grid cell, and the cell was considered a potential KBA if the percentage exceeded the relevant threshold. Criterion B1 refers to geographically restricted species, which are defined as any species whose population is so concentrated that at least 10% of the global population is found within a site (IUCN, 2020). For each species, we calculated the proportion of AOH inside each grid cell, and the cell was considered a potential KBA if the percentage exceeded 10% of the total AOH of the species.

**Table 1.**
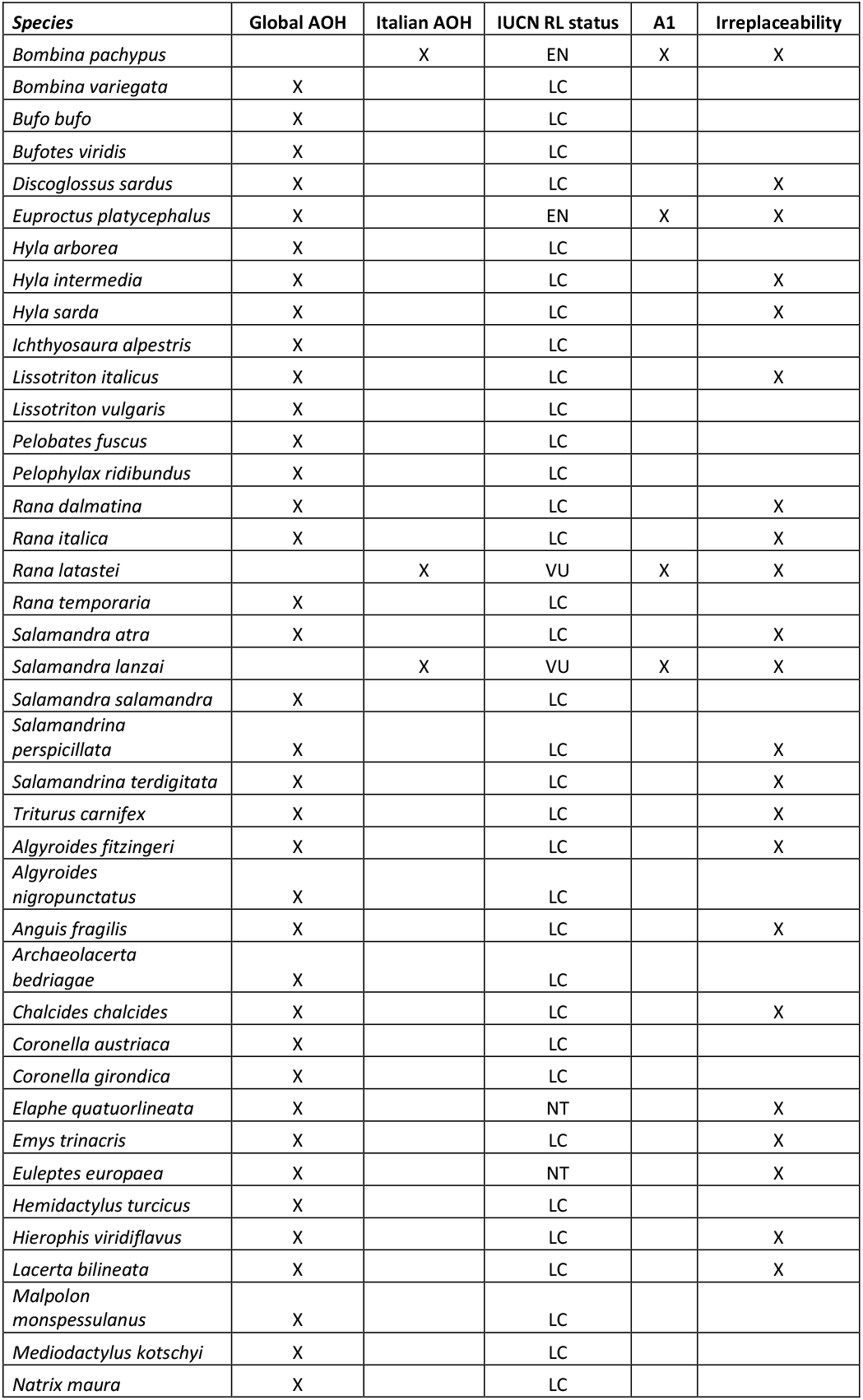
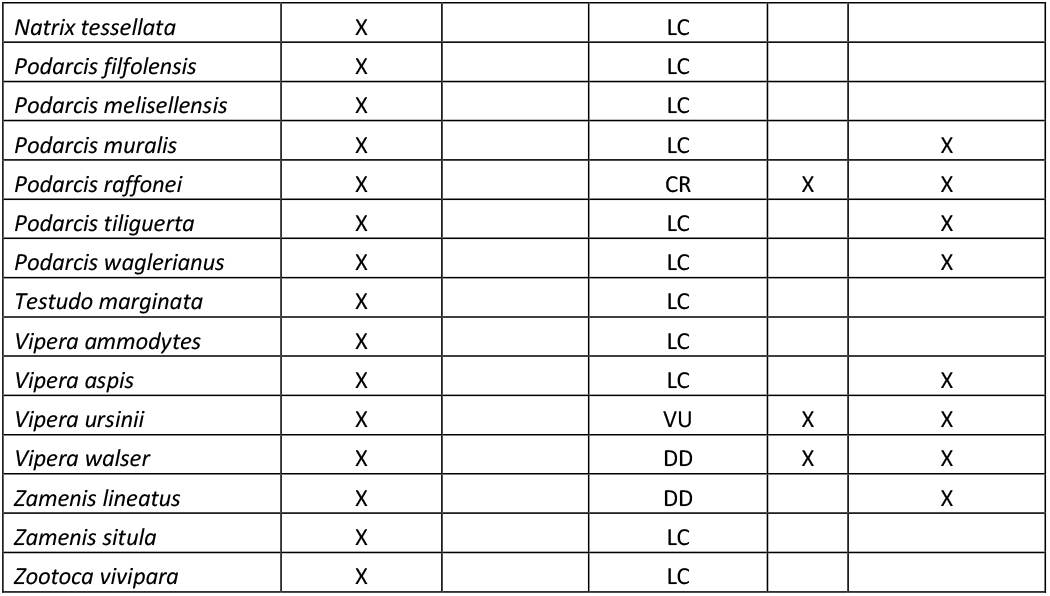
List of the species, type of AOH maps, IUCN status, detailing which species were tested for criteria A1 and for Irreplaceability (criterion E).

Criterion B2 refers to co-occurring geographically restricted species, which are defined as species with a global range size ≤10000 km^2^that co-occur within a site. A site triggers a potential KBA under criterion B2 if it holds ≥1% of the global population of at least 2 geographically restricted species (IUCN, 2020). First, we identified geographically restricted species which have a global range ≤10000 km^2^. For each of these species, we calculated the proportion of AOH inside each grid cell and retained the cells containing more than 10% of the total AOH. Finally, we overlapped these cells across species, and we considered potential KBAs those that were retained for at least 2 species.

Criterion B3 refers to geographically restricted assemblages. The KBA identification guidelines state that: “A site qualifies as potential KBA under criterion B3 if it holds ≥5 species within a taxonomic group or 10% of the species restricted to the ecoregion, whichever is larger” (IUCN 2020). According to the guidelines, a species can be considered ecoregion-restricted if at least 95% of its global population is confined to a single ecoregion. In order to test criterion B3, we used WWF Palaearctic Terrestrial Ecoregion map (Olson et al., 2001), which is freely available at the WWF portal (https://www.worldwildlife.org/publications/terrestrial-ecoregions-of-the-world). Of these maps, we retained only the portion of ecoregions which fell within the administrative boundaries of Italy. We counted a total of 10 ecoregions, listed in Appendix S1. First, we calculated the percentage of global population of the species, based on the AOH maps that was confined to each of the 10 ecoregions. Only species for which at least 95% of the global population is restricted to a single ecoregion were considered candidate species potentially able to trigger criterion B3. All raster grids of ecoregion-restricted species were filtered according to criterion B3 thresholds: only cells hosting ≥0.5% of the global population of the species were retained. Finally, for each resolution, we considered potential KBAs under B3 only areas which matched the B3 population threshold for at least 5 ecoregion-restricted species.

Criterion E refers to the irreplaceability of a site, as measured through quantitative analysis. A site may trigger a potential KBA under criterion E if it has a level of irreplaceability of at least 0.9 on a 0-1 scale (IUCN, 2016). Irreplaceability is the likelihood that the conservation of a given site is needed to achieve a set of conservation targets (Ferrier et al., 2000). We used Marxan’s selection frequency as a proxy to measure irreplaceability (Ball et al., 2009). Given a set of conservation goals, Marxan identifies through a stochastic process a set of spatial planning solutions to achieve the goals. The selection frequency reflects the number of times a specific site has been selected across the set of solutions identified by repeated Marxan runs. According to the KBA guidelines, the target must be set according to the number of mature individuals hosted in a site. Global population size can be inferred through the AOH. If the global range of a species is less than 1,000 km^2^, the whole population should become the target to conserve (IUCN, 2020). The target can be set at 1000 km^2^ if this is the largest value among the other possible values listed in the KBA guidelines (IUCN, 2020). However, since we aimed at identifying KBAs at the national scale, we scaled the targets for species with a global range ≥ 1,000 km^2^ according to the percentage of the global range which is actually found within the boundaries of Italy. Moreover, we did not include in the irreplaceability analysis all species in our dataset for which less than 10% of their global range falls within Italy. In doing so, we avoided potential errors due to marginal fractions of the global range of widely distributed species. All species included in the irreplaceability analysis are reported in Table 1.

Marxan requires a set of planning units to find optimal solutions to meet the conservation targets. We used grid cells as the planning units and overlaid them with AOH maps to quantify the amount of AOH in each cell. Marxan also allows to assign a cost to each planning unit, to be minimized when computing the solutions. We calculated the cost of each planning unit based on the percentage of land within the planning unit. A cost value between 0 (all water) and 1 (all land) was assigned to the planning units. The cost assigned to planning units falling along the coastline reflects the percentage of land area within the planning unit. Marxan analyses were run separately for amphibian and reptile species and repeated for the 16 sets of planning units. The number of runs for each analysis was set to 1000. For each cell resolution, we identified as potential KBAs all cells with selection frequency ≥900.

### Sensitivity and overlap analysis

We tested the response of KBA criteria in relation to the different cell sizes used to detect KBAs. This was done by calculating the total AOH found within the cells identified as potential KBAs for each species at a specific cell resolution, regardless of the position of the grid on the Italian peninsula. The sensitivity analysis was performed for criteria A1 and B1, as they could be triggered by single species. A measure of the degree of overlap between KBAs identified using different criteria was obtained by summing the total AOH identified as KBA for all species under each criterion. Subsequently, we examined the percentage of potential Key Biodiversity Areas identified under each criterion, which was nested in the area of other tested criteria. We followed the same procedure for KBAs identified using each of the four cell resolutions. Criterion E was excluded from the analysis, as the method for measuring irreplaceability does not allow to link the KBA to a single species, and therefore to the AOH which triggered criterion E.

Finally, we calculated the total area identified as potential KBA for all criteria, using each of the four cell resolutions. Thus, for each of the implemented cell resolutions, we produced a map of the total AOH area identified as KBA, regardless of the species or the criteria which triggered it. Subsequently, we measured the percentage of potential KBA falling within the current KBA network, according to the World Database of Key Biodiversity Areas (https://www.keybiodiversityareas.org/). Additionally, we measured the percentage of potential KBA which falls within the Natura 2000 network (EEA, 2021) and the national protected area network of Italy (NNB, 2021).

## Results

### Potential KBAs under main criteria A and B

Potential KBAs were detected for multiple species under different criteria (Figure 2 and 3). Criterion A1 was applied to 4 species of amphibians and 3 species of reptiles, because they hold a ‘Threatened’ status in the IUCN Red List (IUCN, 2021). The total extent of potential KBAs identified for the target species decreased with cell size (Figure 2). While the majority of potential KBAs were identified under a single criterion, some areas triggered multiple KBA criteria regardless of the resolution of the grid cell units (Figure 2).

**Figure 2:**
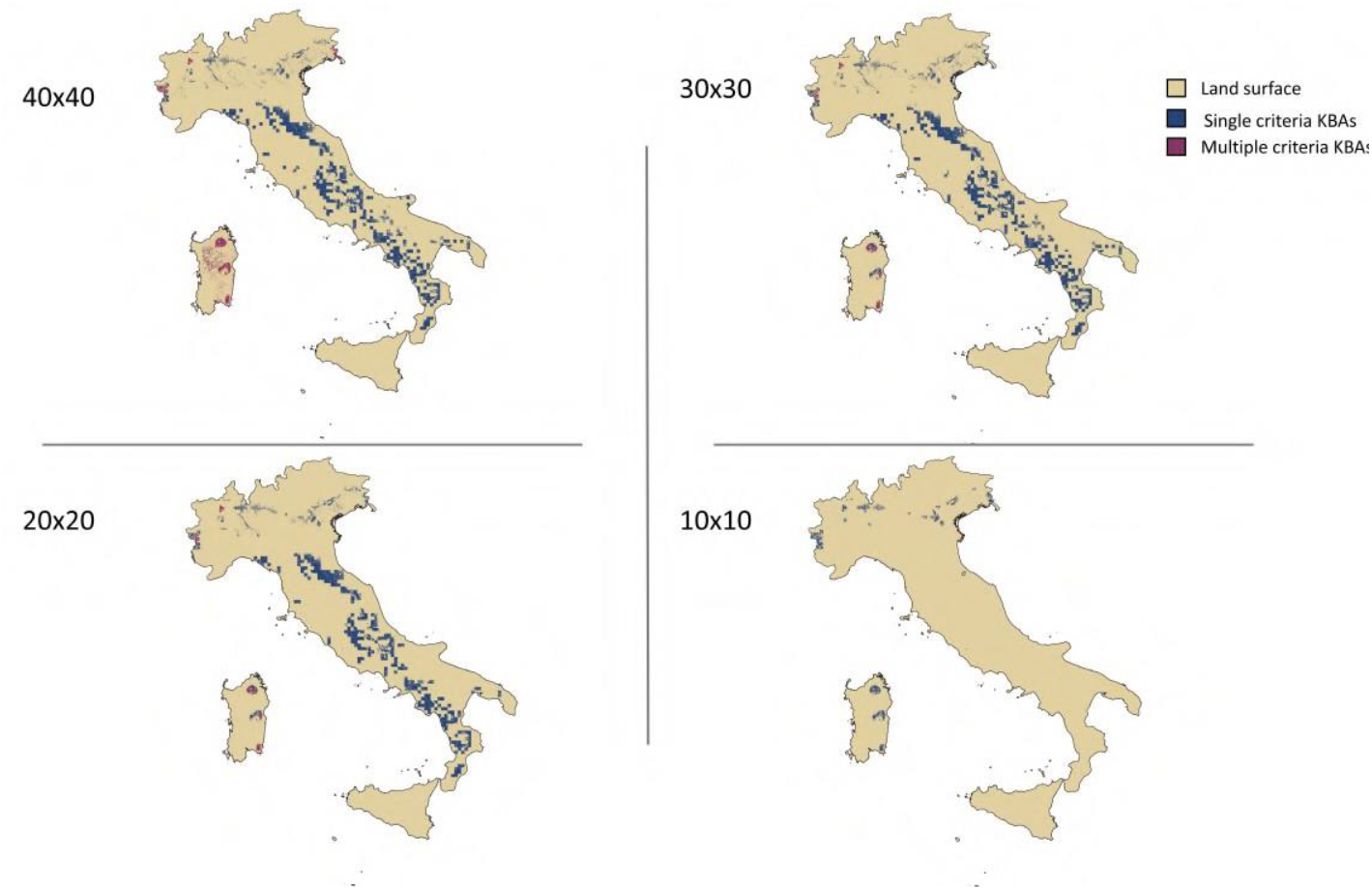
Potential Key Biodiversity Areas identified using all four grid cell resolutions.

**Figure 3:**
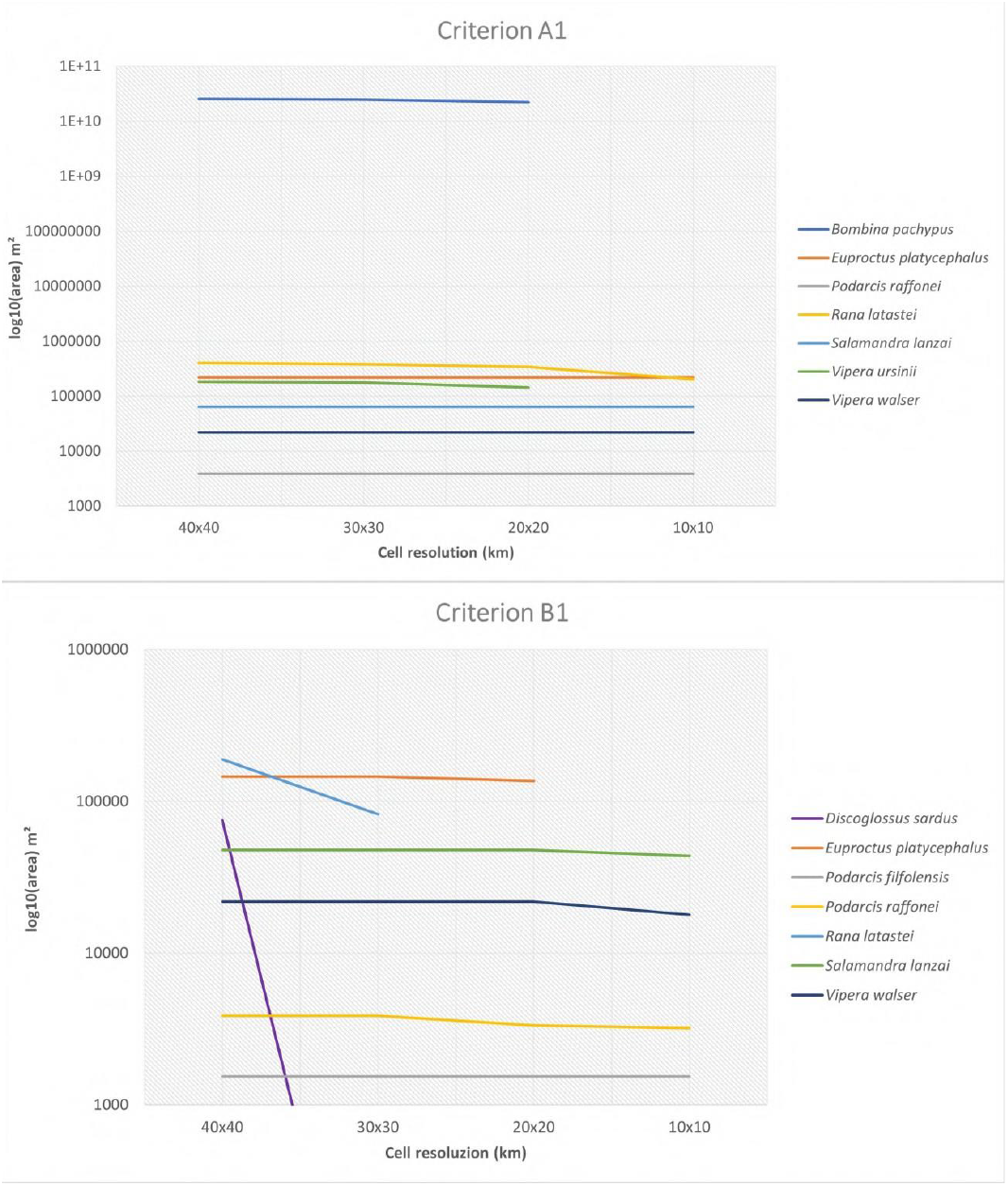
Extent of KBAs under criteria A1 and B1 identified for each species using different grid cell resolutions.

Potential Key Biodiversity Areas under criterion A1 were detected for seven species using 40×40, 30×30 and 20×20 km grid cells, while only five species triggered potential KBAs under criterion A1 using the 10×10 km grid cells, as it is shown in Figure 3. Total area of potential KBA under criterion A1 for *Bombina pachypus* and *Vipera ursinii* dropped to zero when using a 10×10 km cell size. Potential KBAs for *Rana latastei* show a slight decrease of extent with decreasing cell size, with the maximum decrease rate observed when using 10×10 km cell size (Figure 3). For the remaining four species, the potential KBA extent did not vary significantly according to the cell resolution.

We detected potential KBAs under criterion B1 for six species with a grid cell size of 40×40 km. With a resolution of 30×30 km, the number of species which could trigger KBAs under criterion B1 was reduced to four, while only for three species we could trigger B1 using 10×10 km grid resolution. The potential KBA extent of *Discoglossus sardus* and *Rana latastei* responded to the variation of cell size by quickly dropping to zero when using smaller cell sizes. *Euproctus platycephalus* showed a slightly decreased KBA extent using 30×30-20×20 km cells, with potential KBA extent dropping to zero when using 10×10 km cells. The remaining three species showed very little to no variation of potential KBA extent using different cell sizes (Figure 3). Criterion B2 could be triggered only by two species of amphibians, *Discoglossus sardus* and *Euproctus platycephalus*, using 40×40 km grid cell size. The total potential KBA area under B2 was concentrated in the island of Sardinia (Figure 2). Concerning criterion B3, we could not identify any area which could satisfy all requirements of B3 and thus trigger a potential KBA under this criterion.

Percentage of overlap between potential KBAs identified under different criteria is shown in Table 2. Regardless of the implemented cell size, the percentage of potential KBAs detected under criterion A1 which was nested within potential KBAs for B1 or B2 remained below 8%. Criterion B1 was nested within potential KBAs for A1 from 30% to 62% of its extent depending on the implemented cell size. Criterion B2 could be triggered only using 40×40 km cells and showed 27% and 71% overlap with potential KBAs identified under criteria A1 and B2 respectively. The percentage of potential KBAs under criterion A1 nested within potential KBAs for B1 was 7.6% using 10×10 km cells and 3.7% using 40×40 km cells. The same trend was observed for the percentage of potential KBAs for B1 which are nested within potential KBAs for A1 (63% with 10×10 km cells, 31% with 40×40 km cells).

**Table 2:** For each cell resolution, we report the percentage (%) of cells identified by each criterion (left column) that are also identified by other criteria.

|  | A1 | B1 | B2 |
| --- | --- | --- | --- |
| **40x40** |  |  |  |
| **A1** | 100 | 3.7 | 1.86 |
| **B1** | 30.67 | 100 | 39.6 |
| **B2** | 27.47 | 70.78 | 100 |
| **30x30** |  |  |  |
| **A1** | 100 | 3.05 | / |
| **B1** | 39.75 | 100 | / |
| **20x20** |  |  |  |
| **A1** | 100 | 3.32 | / |
| **B1** | 56.56 | 100 | / |
| **10x10** |  |  |  |
| **A1** | 100 | 7.62 | / |
| **B1** | 62.58 | 100 | / |

### Overlap with current KBA, Protected Area, and Natura 2000 networks in Italy

The percentage overlap of the potential KBAs we identified with the existing KBA, Protected Area, and Natura 2000 networks in Italy are shown in Table 3. The new potential KBAs detected with 10×10 km cells in this study showed an overlap of 18% with the current KBA network, while using larger cells the percentage of overlap remained stable around 25%. The lowest percentage of overlap between the new potential KBAs and the Italian national protected areas network was observed when using 10×10 km cells. For potential KBAs identified with larger cells the percentage of overlap was 22% to 24%. The Natura 2000 network enclosed from 33% to 35% of the new potential KBAs identified in this study with 20×20 to 40×40 km cells, respectively, with the lowest percentage of overlap observed with KBAs identified through a 10×10 km cell (28%). The mean percentage of new potential KBAs not encompassed by the current KBA network using the four different cell sizes was equal to 70%, while the mean percentage not included in national designated protected areas was 81%. The Natura 2000 network currently excludes a mean of 68% new KBAs using the four cell resolutions.

**Table 3:** Percentage (%) overlap between candidate KBAs and the current KBA, Protected area and Natura 2000 networks.

|  | current KBAs | National PA | N2000 |
| --- | --- | --- | --- |
| 40x40 | 24.35 | 21.86 | 33.37 |
| 30x30 | 24.48 | 22.91 | 33.99 |
| 20x20 | 25.28 | 24.02 | 35.21 |
| 10x10 | 18.04 | 8.28 | 28.59 |

### Potential KBAs under criterion E (Irreplaceability)

Irreplaceable areas for amphibians are reported on the western Italian alps and in the island of Sardinia (Figure 4). In Sardinia, the extent of irreplaceable area increased when larger planning units were implemented. A single planning unit in the north of the island was irreplaceable using 10×10 cells. When using larger planning units, the irreplaceable area extended towards the central and southern parts of the island. Irreplaceability maps for reptiles showed a high density of irreplaceable planning units in Sicily along with the Eolian islands, as well as in a small area of North-western Italian alps independently of the size of the planning unit (Figure 4). A few small areas in the South-eastern part of the country and in Sardinia were reported as irreplaceable, but these areas were not the same across different planning unit sizes. For all identified irreplaceable areas we provided a list of species of which area of habitat is found within them (Appendix S1).

**Figure 4:**
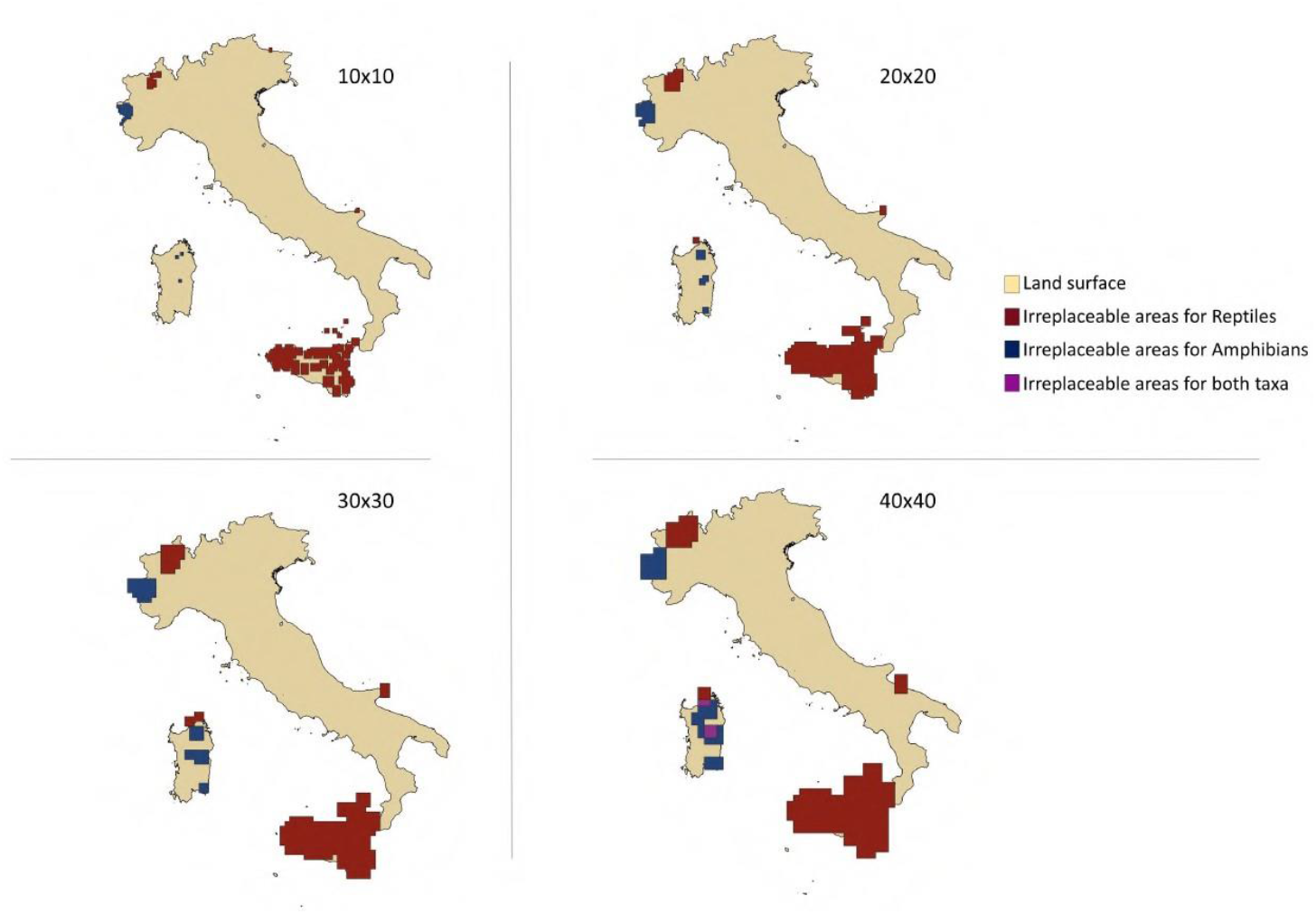
Extent of irreplaceable areas triggered by criterion E for reptiles (red) and amphibians (blue), purple sites are irreplaceable for both taxa.

## Discussion

### Systematic detection of KBAs

Using a set of grids with different cell sizes that can move along the land surface and scan the distribution of species’ habitat availability, allowed us to detect multiple sets of potential KBAs. The number of output potential KBA maps can be high, depending on the number of cell resolutions and grid positions implemented to detect KBAs under different criteria. However, maps of potential KBAs identified using the same cell size can be combined to obtain a single map of KBAs, identified under single or multiple criteria. The resulting map accounts for all identified KBAs, independently of which specific cell and which grid configuration detected it. The extent of potential KBAs identified for criteria A1 and B1, as well as the number of species that were able to trigger them was affected by the size of the cell. For species with a relatively wide distribution range such as *Bombina pachypus* for criterion A1 and *Euproctus platycephalus* for B1, potential KBAs can only be identified using larger cell sizes, as the total KBA extent drops to zero using 10×10 km cells (Figure 3). The opposite is true for microendemics species such as *Vipera walser, Podarcis raffonei* and *Salamandra lanzai*, for which the extent of potential KBA is not affected by the size of the grid cells and remains constant. Thus, the selection of cell size is particularly relevant when testing KBA criteria for species which do not have a very narrow distribution range. Testing different sizes of grid cells provides information on the response of single or multiple species to the KBA criteria in the process of defining new potential KBAs.

We could identify potential KBAs under criterion B2 for 2 species only when using the largest cells (40×40 km), and no potential KBAs under criterion B3 were detected regardless of the implemented cell size. While A1 and B1 criteria can be triggered by a single species, both B2 and B3 criteria depend on the population distribution of more than one species, as they respectively refer to geographically co-occurring species and species assemblage (IUCN, 2020). In the Global Standards for the Identification of Key Biodiversity Areas, an assemblage is defined as a set of species within a taxonomic group for which 95% of the distribution range is confined to an ecoregion or bioregion for at least one life-history stage, or multiple species which share their most important habitats (IUCN, 2020). Despite being the richest Italian region for endemic species of amphibians (Sindaco et al., 2006), our approach could not detect potential KBAs under criterion B3 in the island of Sardinia, as well as in the rest of the peninsula using all four cell resolutions implemented in this study. The minimum cell size able to detect KBAs under criteria B2 and B3 is, as expected, linked to density and proximity of different species populations within a specific area.

Our method was developed to efficiently detect potential KBAs for a set of species under multiple criteria, but not intended to delineate and propose new KBAs, as this process requires different approaches (IUCN, 2020). The purpose of use of this method is to support rapid KBA assessments in different regions of the planet, regardless of their geographic extent. The assessment should be followed by a more accurate evaluation of the identified potential KBA sites to define the true boundaries of a KBA.

### Extent, overlap and nestedness of KBAs

Of all the potential KBAs identified using single criteria, the largest potential KBA network was identified using criterion A1, for which the extent of triggered sites covers, on average using all four cell sizes, 24226 km^2^ of land surface. A large portion of the total area in central Italy for A1 was triggered by *B. pachypus*, an endemic species of toad (Canestrelli et al., 2006). However, the species has been facing population decline over its distribution range in the Apennines during the last two decades (Stagni et al., 2010; Mori & Giovani, 2012; Talarico et al., 2020). Thus, many of the identified potential KBAs for criterion A1 may include sites where the species is not present anymore or close to local extinction.

Systematic conservation planning practices have been compared to the KBA identification process, as the two approaches share broad similarities in the way they aim at identifying important sites for biodiversity (Smith et al., 2018). In the context of conservation planning, using smaller planning units to achieve specific conservation targets was proven to be more efficient than using larger planning units (Pressey & Logan, 1998). The use of smaller planning units tends to maximize spatial and cost efficiency leading to easier compromises, and therefore provides better solutions to design conservation networks (Pressey & Logan, 1998; Hamel et al., 2012; Cheok et al., 2016). The percentage of overlap between potential KBAs detected using different criteria showed a tendency to decrease when using larger cells (Table 2). Thus, keeping the cell unit small may limit the KBA detection to areas which are most relevant for the conservation of one or several species under multiple criteria.

The overlap scores between the total network of potential KBAs (A1, B1 and B2) and current KBAs, national protected areas and Natura 2000 sites indicate that, regardless of the implemented cell size, a high percentage of the identified potential KBAs is currently located outside of the three different networks of important biodiversity areas (Table 3). In particular, ≥75% is excluded from the current KBA network, ≥76% is excluded from national designated protected areas and ≥65% is not located within the Natura 2000 network. We stress the importance of considering the identified sites as potential KBAs and not as proposed KBAs, as the presence of the species in those sites must be confirmed with a minimum threshold of mature individuals (IUCN, 2020). Therefore, the effective potential KBA extent of a site may be reduced when undergoing the proposal process. However, the current KBA network in Italy proves to be insufficient at capturing potential KBAs for non-avian taxa. As new potential KBAs will be detected for other species, this evidence is likely to become stronger. Thus, rapid assessments of potential KBAs for high numbers of species will play an important role in bridging the gap of knowledge on the true distribution of KBAs globally. Methods to systematically apply KBA criteria such as the one presented in this study, can significantly accelerate the KBA mapping process.

### Irreplaceable sites

Areas with a high value of irreplaceability for both amphibians and reptiles were detected mainly, but not exclusively, where endemic species are known to occur (Tessa et al., 2007; Ficetola et al., 2021; Ficetola et al., 2020; Salvi et al., 2017). This suggests that the endemic status of a species is an important factor affecting the identification of KBAs under criterion E. Areas hosting endemic and microendemic species are more likely to be identified as irreplaceable. Using larger cells as planning units for our irreplaceability analysis implied the identification of larger irreplaceable sites which most likely do not represent important sites for the species which triggered criterion E. A large cell unit would capture a sufficient percentage of the global habitat distribution of a species to trigger criterion E. However, it also tends to capture a large amount of area where the species does not occur, simply because the whole planning unit will result as irreplaceable and thus, equally contributing to the achievement of the conservation targets. This circumstance is particularly evident in the areas hosting microendemic species, such as *Salamandra lanzai, Podarcis raffonei* and *Vipera walser*. The implementation of criterion E identified important areas of biodiversity which were not detected by criteria A1, B1, B2 and B3. Thus, underlying the important role of Criterion E as a support to the other KBA criteria, as previously suggested (Di Marco et al., 2016; Smith et al., 2018). For instance, a large extent of irreplaceable sites was identified within the island of Sicily, although these sites did not trigger KBAs under other criteria. This suggests that an integration of KBA criteria is needed to perform large scale assessments of potential KBAs.

## Supporting information

Appenidx 2

