## Appendix 1 for "An automated approach for systematic detection of Key Biodiversity Areas"

**Additional Area of Habita (AOH) maps**

Additional Area of habitat (AOH) maps for three species (Table 1) endemic to Italy were produced based on global distribution ranges obtained from the from the Atlas of Amphibians and Reptiles of Italy, curated by the Societas Herpetologica Italica (SHI). The maps are available upon request from SHI. Species were linked to their natural habitat using Copernicus Global Land Service Land Cover (CGLS-LC100) 2019 classes as a habitat surrogate. CGLS-LC100 is a global map of land cover produced by the European Union's Earth observation programme, freely accessible at the official programme website (land.copernicus.eu). The map is available in a standard latitude/longitude grid (EPSG:4326), ellipsoid WGS 1984. The grid resolution is 1°/1008. Information on land cover categories representing habitat for the species, as well as altitude limits information was obtained from Nania et al. (2022). The CGLS-LC100 map was combined with altitude information from the Shuttle Radar Topography Mission (USGS EROS Archive, 2019) map, which has a resolution of approximately 30 m and was resampled to match the CGLS-LC100 resolution. to produce a base map. Each raster cell in the base map holds information on the land cover class and altitude data. We used SHI distribution ranges to mask the area of the base map which is found out of the species range. Then, the base map was filtered according to species habitat requirements to produce the area of habitat maps.

**Irreplaceable areas identified with four different cell resolutions and the list of species found in each of the Irreplaceable regions**

For each numbered irreplaceable area identified with the four cell sizes, a list of species occurring in the area is provided. Areas in red have been identified as irreplaceable for reptile species. Areas in blue are identified as irreplaceable for amphibian species. Purple areas were identified as irreplaceable for both taxa.

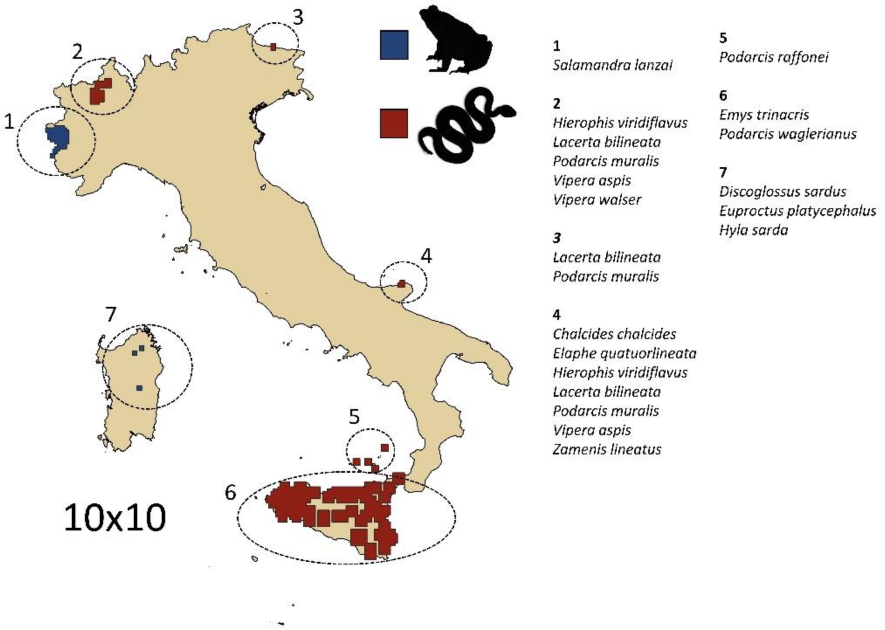

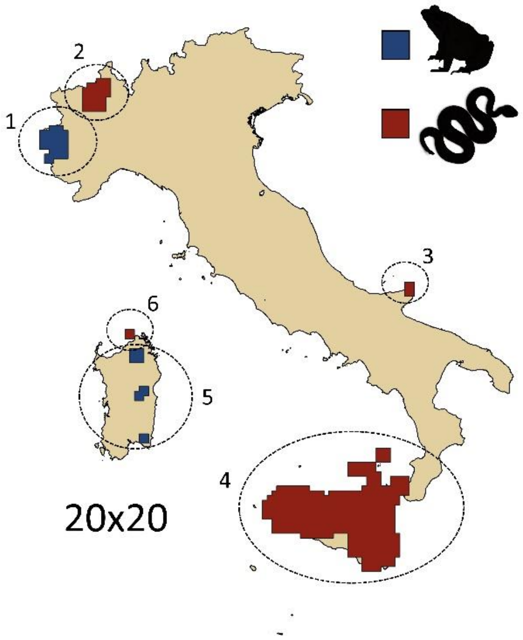


**1**
*Salamandra lanzai*
*Triturus carnifex*

**2**
*Hierophis viridiflavus*
*Lacerta bilineata*
*Podarcis muralis*
*Vipera aspis*
*Vipera walser*

***3***
*Chalcides chalcides*
*Hierophis viridiflavus*
*Lacerta bilineata*
*Podarcis muralis*
*Vipera aspis*
*Zamenis lineatus*

**4**
*Emys trinacris*
*Podarcis waglerianus*
*Podarcis raffonei*

**5**
*Discoglossus sardus*
*Euproctus platycephalus*
*Hyla sarda*

**6**
*Algyroides fitzingeri*
*Archaeolacerta bedriagae*
*Chalcides chalcides*
*Euleptes europaea*
*Hierophis viridiflavus*
*Podarcis tiliguerta*

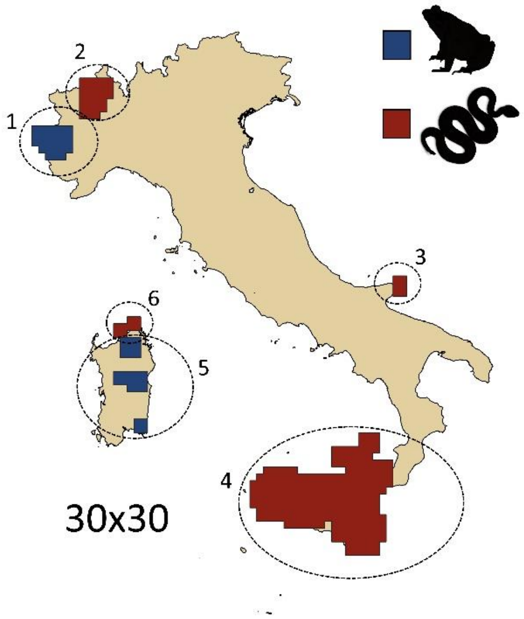


**1**
*Rana latastei*
*Salamandra lanzai*
*Triturus carnifex*

**2**
*Hierophis viridiflavus*
*Lacerta bilineata*
*Podarcis muralis*
*Vipera aspis*
*Vipera walser*

***3***
*Chalcides chalcides*
*Elaphe quatuorlineata*
*Hierophis viridiflavus*
*Lacerta bilineata*
*Podarcis muralis*
*Vipera aspis*
*Zamenis lineatus*

**4**
*Emys trinacris*
*Podarcis waglerianus*
*Podarcis raffonei*

**5**
*Discoglossus sardus*
*Euproctus platycephalus*
*Hyla sarda*

**6**
*Algyroides fitzingeri*
*Archaeolacerta bedriagae*
*Chalcides chalcides*
*Euleptes europaea*
*Hierophis viridiflavus*
*Podarcis tiliguerta*

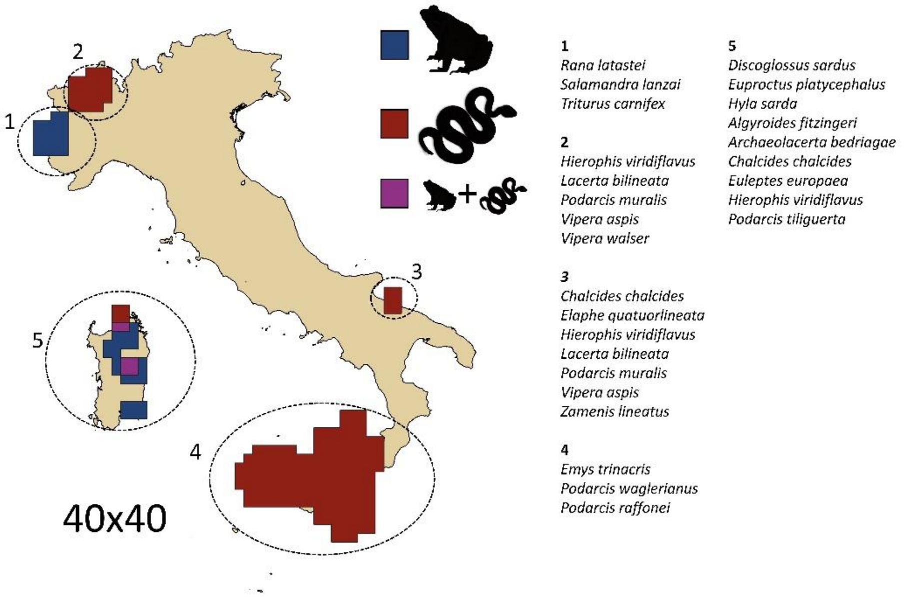

1
2
3
4
5
40x40
1
Rana latastei
Salamandra lanzai
Triturus carnifex
2
Hierophis viridiflavus
Lacerta bilineata
Podarcis muralis
Vipera aspis
Vipera walser
3
Chalcides chalcides
Elaphe quatuorlineata
Hierophis viridiflavus
Lacerta bilineata
Podarcis muralis
Vipera aspis
Zamenis lineatus
4
Emys trinacris
Podarcis waglerianus
Podarcis raffonei
5
Discoglossus sardus
Euproctus platycephalus
Hyla sarda
Algyroides fitzingeri
Archaeolacerta bedriagae
Chalcides chalcides
Euleptes europaea
Hierophis viridiflavus
Podarcis tiliguerta
