## Supplementary material for "An automated approach for systematic detection of Key Biodiversity Areas": Appenidx 2

**GRASS and R codes used to build AOH maps and perform the KBA assessment**

```
##################################################################
### GRASS CODE TO GENERATE AOH MAPS ###

# Create list of files

cd /media/reclass

ls > hlist.txt
#import range
for i in `cat /media/hlist.txt`
do
v.in.ogr input=/media/$i.shp output=$(echo $i | sed 's/.shp *//')
snap=1e-09 --overwrite
done


#Rasterize


for i in `cat /media/nas/userdata/hlist.txt`
do
g.region -a vector=$i res=0:00:03.571
v.to.rast input=$i type=area use=val output=ras_$i --overwrite
r.mapcalc "$i = $i" --overwrite
done


# produce AOH
for i in `cat /media/nas/userdata/hlist.txt`
do
g.region -a raster=ras_$i
r.mask raster=ras_$i --overwrite

r.reclass in=bm_CGLS out=$i rules=/media/reclas/$i --overwrite r.colors map=$i rules=-<<EOF
0  160:160:160
1 5:110:7
EOF
r.mapcalc "$i = $i" –overwrite
r.mask -r
done


########################################################
```

```
####################################################################################################
## CODE TO SCAN THE GEOGRAPHIC SURFACE AND CAPTURE HABITAT EXTENT WITHIN CELLS

# 10 KM

####################################################################################################
##

# start GRASS session

grass /data/user/grassdata/equal_area_eu/maps

# import raster grid

r.import input=/media/nas/userdata/user/raster.tif output=my_grid --overwrite

# import AOH

for i in `cat /media/nas/userdata/user/specieslist.txt`

do

r.in.gdal input=/media/nas/userdata/user/species_raster/$i'.tif' output=$(echo $i) --overwrite

done

# moving grid

for i in `cat /media/nas/userdata/user/specieslist.txt`

do

cd /media/nas/userdata/user/moving_grid_res

mkdir $(echo $i)

cd /media/nas/userdata/user/moving_grid_res/$(echo $i)

g.region raster=my_grid -p

g.region res=10000 -ap

r.resamp.stats -w input=$(echo $i) output=$(echo $i)_standard method=sum --overwrite

r.out.gdal input=$(echo $i)_standard output=$(echo $i)_standard'.tif'

#set the region

# reset region: move X

g.region raster=my_grid n=2710000 s=1380000 w=4015000 e=5115000 -p

r.resamp.stats -w input=$(echo $i) output=$(echo $i)_E method=sum --overwrite

r.out.gdal input=$(echo $i)_E output=$(echo $i)_E'.tif'


# reset region: move Y

g.region raster=my_grid n=2715000 s=1385000 w=4010000 e=5110000 -p

r.resamp.stats -w input=$(echo $i) output=$(echo $i)_N method=sum --overwrite

r.out.gdal input=$(echo $i)_N output=$(echo $i)_N'.tif'


# reset region: move diagonal

g.region raster=my_grid n=2715000 s=1385000 w=4015000 e=5115000 -p

r.resamp.stats -w input=$(echo $i) output=$(echo $i)_NE method=sum --overwrite
```

```
r.out.gdal input=$(echo $i)_NE output=$(echo $i)_NE'.tif'

done

####################################################################################################
##
# 20 KM
####################################################################################################
##
# moving grid
for i in `cat /media/nas/userdata/user/specieslist.txt`
do
cd /media/nas/userdata/user/moving_grid_res20
mkdir $(echo $i)
cd /media/nas/userdata/user/moving_grid_res20/$(echo $i)

g.region raster=my_grid -p
g.region res=20000 -ap
r.resamp.stats -w input=$(echo $i) output=$(echo $i)_standard method=sum --overwrite
r.out.gdal input=$(echo $i)_standard output=$(echo $i)_standard'.tif'
#now the region is set as n=2720000 s=1380000 w=4000000 e=5120000

# reset region: move 20Km X
g.region n=2720000 s=1380000 w=4010000 e=5130000  res=20000 -p
r.resamp.stats -w input=$(echo $i) output=$(echo $i)_E method=sum --overwrite
r.out.gdal input=$(echo $i)_E output=$(echo $i)_E'.tif'

# reset region: move Y
g.region n=2730000 s=1390000 w=4000000 e=5120000  res=20000 -p
r.resamp.stats -w input=$(echo $i) output=$(echo $i)_N method=sum --overwrite
r.out.gdal input=$(echo $i)_N output=$(echo $i)_N'.tif'

# reset region: move diagonal
g.region n=2730000 s=1390000 w=4010000 e=5130000  res=20000 -p
r.resamp.stats -w input=$(echo $i) output=$(echo $i)_NE method=sum --overwrite
r.out.gdal input=$(echo $i)_NE output=$(echo $i)_NE'.tif'

done



####################################################################################################
##
```

```
# 30 KM
####################################################################################################
##

# moving grid
for i in `cat /media/nas/userdata/user/specieslist.txt`
do
cd /media/nas/userdata/user/moving_grid_res30
mkdir $(echo $i)
cd /media/nas/userdata/user/moving_grid_res30N/$(echo $i)
g.region raster=my_grid -p
g.region res=30000 -ap
r.resamp.stats -w input=$(echo $i) output=$(echo $i)_standard method=sum --overwrite
r.out.gdal input=$(echo $i)_standard output=$(echo $i)_standard'.tif'
#now the region is set as n=2730000 s=1380000 w=3990000 e=5130000

# reset region: move X
g.region n=2730000 s=1380000 w=4005000 e=5145000  res=30000 -p
r.resamp.stats -w input=$(echo $i) output=$(echo $i)_E method=sum --overwrite
r.out.gdal input=$(echo $i)_E output=$(echo $i)_E'.tif'
# reset region: move Y
g.region n=2745000 s=1395000 w=3990000 e=5130000  res=30000 -p
r.resamp.stats -w input=$(echo $i) output=$(echo $i)_N method=sum --overwrite
r.out.gdal input=$(echo $i)_N output=$(echo $i)_N'.tif'

# reset region: move diagonal
g.region n=2745000 s=1395000 w=4005000 e=5145000  res=30000 -p
r.resamp.stats -w input=$(echo $i) output=$(echo $i)_NE method=sum --overwrite
r.out.gdal input=$(echo $i)_NE output=$(echo $i)_NE'.tif'

done


####################################################################################################
##
#40 KM
####################################################################################################
# moving grid
for i in `cat /media/nas/userdata/user/specieslist.txt`
do
cd /media/nas/userdata/user/moving_grid_res40
```

```
mkdir $(echo $i)

cd /media/nas/userdata/user/moving_grid_res40/$(echo $i)

#import raster grid and species AOH

g.region raster=my_grid -p

g.region res=40000 -ap

r.resamp.stats -w input=$(echo $i) output=$(echo $i)_standard method=sum --overwrite

r.out.gdal input=$(echo $i)_standard output=$(echo $i)_standard'.tif'

#now the region is set as n=2720000 s=1360000 w=4000000 e=5120000


# reset region: move X

g.region n=2720000 s=1360000 w=4020000 e=5140000  res=40000 -p

r.resamp.stats -w input=$(echo $i) output=$(echo $i)_E method=sum --overwrite

r.out.gdal input=$(echo $i)_E output=$(echo $i)_E'.tif'


# reset region: move Y

g.region n=2740000 s=1380000 w=4000000 e=5120000  res=40000 -p

r.resamp.stats -w input=$(echo $i) output=$(echo $i)_N method=sum --overwrite

r.out.gdal input=$(echo $i)_N output=$(echo $i)_N'.tif'


# reset region: move diagonal

g.region n=2740000 s=1380000 w=4020000 e=5140000  res=40000 -p

r.resamp.stats -w input=$(echo $i) output=$(echo $i)_NE method=sum --overwrite

r.out.gdal input=$(echo $i)_NE output=$(echo $i)_NE'.tif'


done
```

**# R code to apply the KBA criteria**

```
####################################################################################################

# Criterion A1

#############################################################################################################


library(raster)

library(sp)

library(rgdal)

library(icesTAF) # for mkdir


# species list

species_list<-read.csv("/home/SPS.csv", header = F)

KBA_size<-list('10')
```

```
pixel_area<-8187.1200316976 # pixel area in m2

for (row in 1:nrow(species_list)) {
  current_sp<-as.character(species_list[row,1])
  for (j in KBA_size) {
    current_KBA_size<-j
    mkdir(paste("/home/A1_results_", j, "/",current_sp, sep = ""))

    # read area stats of the species
    area_table<-read.csv(paste("/home/area_specs/",
                               current_sp,"_stats.csv",sep = ""), header = F, sep=",")
    total_area<-area_table[2,2]

    # set threshold %

    if (species_list[row,2]=='EN') {
      trsh_num<-0.5
    } else if (species_list[row,2]=='CR'){
      trsh_num<-0.5
    } else { trsh_num<-1 }

    my_trsh<-(total_area/100)*trsh_num # value of threshold % of total AOH area
    # pixel threshold
    pixel_trsh<-(1/pixel_area)*my_trsh



    files <-as.list(list.files(path=paste("/home/moving_grid_res/",
                                          current_sp,"/", sep = ""), pattern = ".tif"))

    for(i in files){
      sliding_map<-raster(paste("/home/",
                                current_sp,"/", i, sep = "")) # need sliding grid maps

      sliding_map[sliding_map[] < pixel_trsh] = NA

      if(!is.na(minValue(sliding_map))){
        writeRaster(sliding_map,
                    filename=paste("/home/A1_results_",
                                   j, "/",current_sp, "/", i,sep = ""), format="GTiff",
overwrite=TRUE)
```

```
      }
    }
  }
}


####################################################################################################
###########
# Criterion B1
####################################################################################################
###########


library(raster)
library(sp)
library(rgdal)
library(icesTAF) # for mkdir

# species list
species_list<-read.csv("/media/hlist.txt", header = F)
KBA_size<-list('10', '20', '30', '40')

pixel_area<-8187.1200316976

for (row in 1:nrow(species_list)) {
  current_sp<-as.character(species_list[row,1])
  for (j in KBA_size) {
    current_KBA_size<-j
    mkdir(paste("/media/B1_results_", j, "/",current_sp, sep = ""))

    # read area stats of the species

    area_table<-read.csv(paste("/media/",
                           current_sp,"_area_stats_B1.csv",sep = ""), header = F, sep=",")
    total_area<-area_table[2,2]

    # set threshold %
    trsh_num<-10

    my_trsh<-(total_area/100)*trsh_num

    # pixel threshold
```

```
    pixel_trsh<-(1/pixel_area)*my_trsh



    files <-as.list(list.files(path=paste("/media/", j, "_moving_grid_res/",
                                          current_sp,"/", sep = ""), pattern = ".tif"))



    for(i in files){
      sliding_map<-raster(paste("/media/nas/userdata/", j, "_moving_grid_res_IUCN/",
                                current_sp,"/", i, sep = "")) # need sliding grid maps

      sliding_map[sliding_map[] < pixel_trsh] = NA # check the use of braces to acces values of the
raster

      if(!is.na(minValue(sliding_map))){
        writeRaster(sliding_map,
                    filename=paste("/media/nas/B1_results_",
                                   j, "/",current_sp, "/", i,sep = ""), format="GTiff",
overwrite=TRUE)
      }
    }
  }
}



##############################################################################################
###########
# Criterion B2
##############################################################################################
###########

library(raster)
library(sp)
library(rgdal)
library(icesTAF) # for mkdir



# species list
species_list<-read.csv("/media/hlist.txt", header = F)
KBA_size<-list('10', '20', '30', '40')
```

```
pixel_area<-8187.1200316976

for (row in 1:nrow(species_list)) {
  current_sp<-as.character(species_list[row,1])
  for (j in KBA_size) {
    current_KBA_size<-j
    mkdir(paste("/media/B2_results_", j, "/",current_sp, sep = ""))

    # read area stats of the species
    area_table<-read.csv(paste("/media/area_specs/",
                               current_sp,"_area_stats.csv",sep = ""), header = F, sep=",")
    total_area<-area_table[2,2]

    # set threshold
    my_area_trsh<-10000000000

    if(total_area<=my_area_trsh){

      my_thresh<-(total_area/100)*1 # value of 1% of total AOH area

      # pixel threshold equivalent to 1% of AOH
      pixel_trsh<-(1/pixel_area)*my_thresh



      files <-as.list(list.files(path=paste("/media/", j, "_moving_grid_res/",
                                            current_sp,"/", sep = ""), pattern = ".tif"))

      for(i in files){
        sliding_map<-raster(paste("/media/", j, "_moving_grid_res/",
                                  current_sp,"/", i, sep = ""))

        sliding_map[sliding_map[] < pixel_trsh] = NA # check the use of braces to acces values of
the raster
        if(!is.na(minValue(sliding_map))){
          writeRaster(sliding_map,
                      filename=paste("/media/B2_results_",
                                     j, "/",current_sp, "/", i,sep = ""), format="GTiff",
overwrite=TRUE)
        }
```

```
      }
    }
  }
}



########################################################################################
# Criterion B3
########################################################################################

setwd("/media/Ecoregion_WWF/")

species_list<-read.csv("/media/sp.txt", header = F)
ECOregions<-read.csv("/media/ecoregions.csv", header = F)

for (row in 1:nrow(species_list)){
  current_sp<-as.character(species_list[row,1])
  # read area stats of the species
  area_table<-read.csv(paste("/media/area_specs/",
                             current_sp,"_area_stats_B1.csv",sep = ""), header = F, sep=",")
  total_area<-as.numeric(area_table[2,2])
  my_thr<-(total_area/100)*95
  for (row in 1:nrow(ECOregions)){
    current_eco<-as.character(ECOregions[row,2])
    ecode<-as.character(ECOregions[row,1])

    if (file.exists(paste("/media/SpxECO/",
      current_sp,"_", ecode, ".csv", sep = ""))) {
        current_ecoarea<-read.csv(paste("/media/Ecoregion_WWF/SpxECO/",
                             current_sp,"_", ecode, ".csv", sep = ""), header = F, sep=",")
        total_ecoarea<-as.numeric(current_ecoarea[2,2])
        if(is.na(total_ecoarea)) {
            next
        } else if (total_ecoarea>=my_thr){
          write(paste(current_sp,
            "\nEcoregion: ", current_eco,
            "\ntotal area: ", total_area,
            "\nTotal area in ecoregion: ", total_ecoarea,
            "\nThreshold", my_thr,
```

```
            "\n\n-----------------"), file="B3_res_amphibians2.txt", append = T)
        } else { print(paste(current_sp," in ", current_eco, " does not trigger B3", sep = ""))}


    } else { print(paste(current_sp,"_", ecode, ".csv"," unavailable", sep = ""))}
  }
}


####################################################################################################
#########################
```
